# CRISPR-SURF: Discovering regulatory elements by deconvolution of CRISPR tiling screen data

**DOI:** 10.1101/345850

**Authors:** Jonathan Y. Hsu, Charles P. Fulco, Mitchel A. Cole, Matthew C. Canver, Danilo Pellin, Falak Sher, Rick Farouni, Kendell Clement, James A. Guo, Luca Biasco, Stuart H. Orkin, Jesse M. Engreitz, Eric S. Lander, J. Keith Joung, Daniel E. Bauer, Luca Pinello

## Abstract

Tiling screens using CRISPR-Cas technologies provide a powerful approach to map regulatory elements to phenotypes of interest, but computational methods that effectively model these experimental approaches for different CRISPR technologies are not readily available. Here we present CRISPR-SURF, a deconvolution framework to identify functional regulatory regions in the genome from data generated by CRISPR-Cas nuclease, CRISPR interference (CRISPRi), or CRISPR activation (CRISPRa) tiling screens. We validated CRISPR-SURF on previously published and new data, identifying both experimentally validated and new potential regulatory elements. With CRISPR tiling screens now being increasingly used to elucidate the regulatory architecture of the non-coding genome, CRISPRSURF provides a generalizable and accessible solution for the discovery of regulatory elements.

## Main Text

The advent of programmable genome editing using CRISPR-based technologies has enabled high-throughput functional interrogation of non-coding elements in the genome^1^^-^^6^. Functional mapping can be achieved by densely tiling single guide RNAs (sgRNAs) across a non-coding region of interest, with each sgRNA linking a unique genomic sequence to an observable phenotype. These tiled sgRNAs can be used with CRISPR-Cas genome-editing nucleases^7,8^, CRISPR interference^9^ (CRISPRi), or CRISPR activation^10^ (CRISPRa) to introduce indel mutations, repress a target element, or activate a target element, respectively. The ability to perform CRISPR tiling screens allows for the systematic and quantitative assessment of causal links between non-coding regulatory elements and biological phenotypes of interest, highlighting its substantial potential to elucidate the regulatory architecture underlying gene expression.

Although experimental protocols are readily available for tiling a genomic region with genetic or epigenetic perturbations, analyzing the resulting sequencing data can be challenging. The computational challenges include contending with (i) the variability in sgRNA targeting efficiencies, (ii) the non-uniform spacing of sgRNAs due to PAM requirements, (iii) the extent of shared information amongst neighboring sgRNAs, and (iv) different perturbations induced by the various CRISPR technologies. We developed CRISPR-SURF (Screening Uncharacterized Region Function) to address these challenges and to provide an open-source command line tool and interactive web-based application for the analysis of CRISPR tiling screen data.

The methodology underlying the CRISPR-SURF framework leverages the concept that sgRNAs represent a functional read-out for base pairs (bp) within its perturbation range. This range has a characteristic size profile that depends on the CRISPR screening approach used: CRISPR-Cas nucleases introduce insertion and deletion (indel) mutations of varying lengths (typically 1-10 bp, although potentially varying with cell type) whereas CRISPRi and CRISPRa strategies may remodel chromatin structure across hundreds of nucleotides. Importantly, each CRISPR technology offers its own advantage: CRISPRi and CRISPRa strategies increase the likelihood of detecting regulatory elements given their larger perturbation ranges, whereas CRISPR-Cas nuclease tiling screens provide higher resolution on the boundaries of regulatory elements given its sharper perturbation window. Because each sgRNA perturbs variable-size regions around its target site, the sgRNA data from CRISPR tiling screens can be seen as imprecise measurements of an underlying genomic regulatory signal. To address this variable, we modeled the generation of these imprecise measurements by means of a convolution operation that accounts for the perturbation profile associated with the CRISPR technology used. CRISPR-SURF deconvolves the CRISPR tiling screen data to infer the underlying genomic regulatory signal **(Figure 1a-b)**.

**Figure 1.**
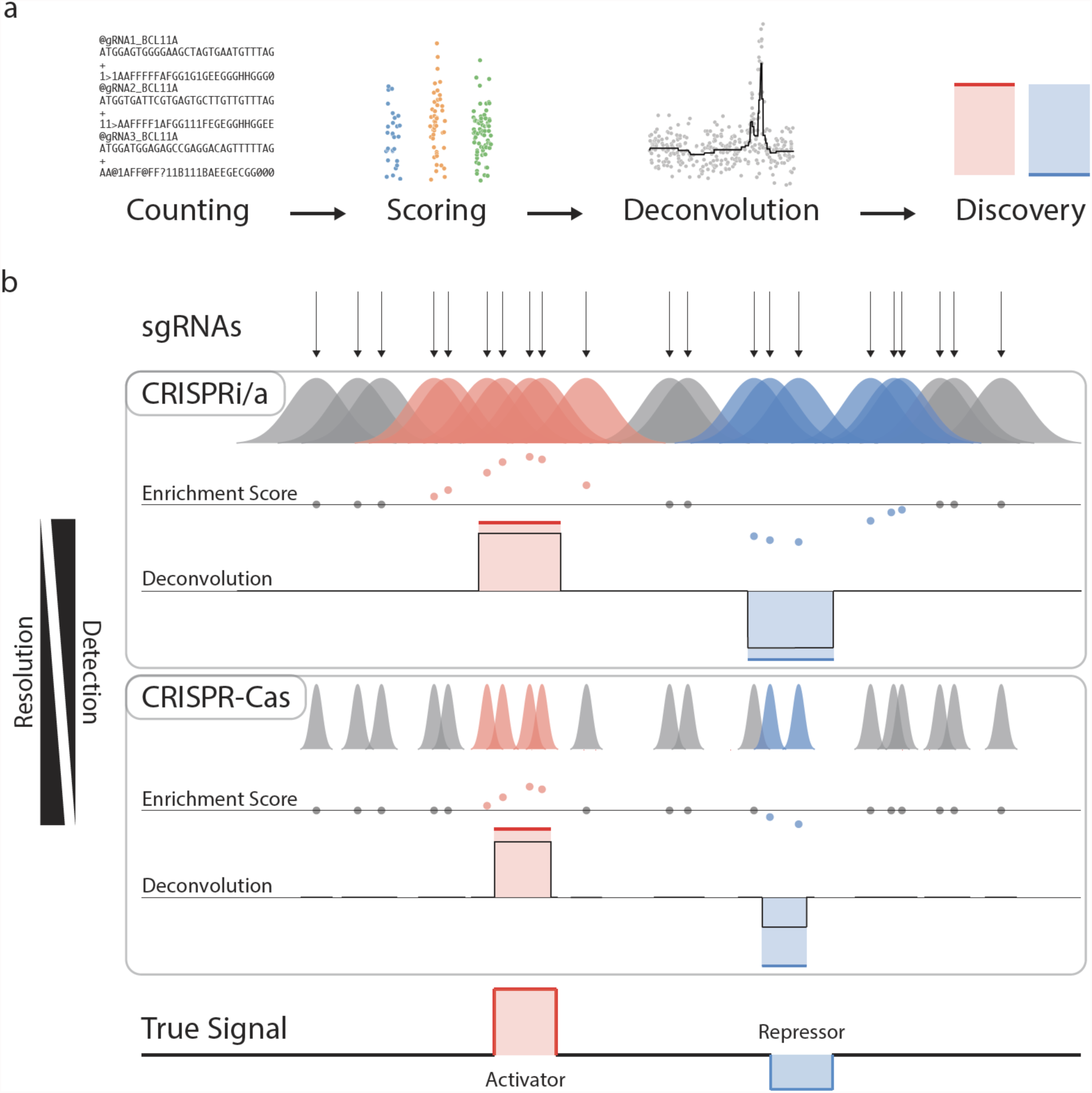
CRISPR-SURF deconvolution framework. **(a)** An overview of the CRISPR-SURF computational pipeline. **(b)** An illustration of the CRISPR-SURF deconvolution framework. Based on sgRNA targeting positions, expected perturbation profiles (CRISPRi/a or CRISPR-Cas), and enrichment scores, the deconvolution algorithm aims to construct a signal that represents the underlying functional signal of the genome (bottom track).

CRISPR-SURF finds the genomic regulatory signal at single bp resolution that best explains the observed sgRNA scores given the perturbation profile and sgRNA spacing. The CRISPR-SURF framework accounts for overlapping perturbation profiles between neighboring sgRNAs and leverages this shared information to infer an underlying genomic regulatory signal even from noisy measurements. The exact sgRNA targeting coordinates are also taken into account, allowing for location-dependent statistical tests that are powered based on the local placement of sgRNAs in a region. This enables CRISPR-SURF to estimate perturbation- and position-specific statistical power for CRISPR tiling screens **(Supplementary Note 1)**.

We evaluated the performance of CRISPR-SURF using three published CRISPR tiling screens spanning CRISPR-Cas9, CRISPRi, and CRISPRa modalities^1,2,3^. For all three data sets, CRISPR-SURF reliably identified all of the experimentally validated regulatory elements, despite the different experimental and computational approaches used in each of these earlier studies. CRISPR-SURF further identified new potentially novel regulatory regions that are also supported by both chromatin accessibility and epigenetic marks **(Supplementary Figure 1– 3, Supplementary Note 2)**. Additionally, we used these validated screens to establish guidelines for the design of more efficient and cost-effective CRISPR tiling screens through down-sampling simulations of the sgRNA libraries **(Supplementary Figure 4, Supplementary Note 3)**.

Next, we looked to characterize CRISPR-SURF performance through a direct comparison of tiling screens using different modalities all targeted to the same locus. We performed two matched CRISPR tiling screens using the *Streptococcus pyogenes* Cas9 nuclease (SpCas9) and CRISPRi (dCas9-KRAB) on the *BCL11A* locus, which encodes a potent transcriptional repressor of fetal haemoglobin (HbF) **(Supplementary Note 4)**. Both tiling screens spanned 55 DNase I hypersensitive sites (DHSs) around the *BCL11A* gene to comprehensively assess the potential role of each of these regions in regulating *BCL11A* **(Figure 2a, Supplementary Figure 7)**.

**Figure 2.**
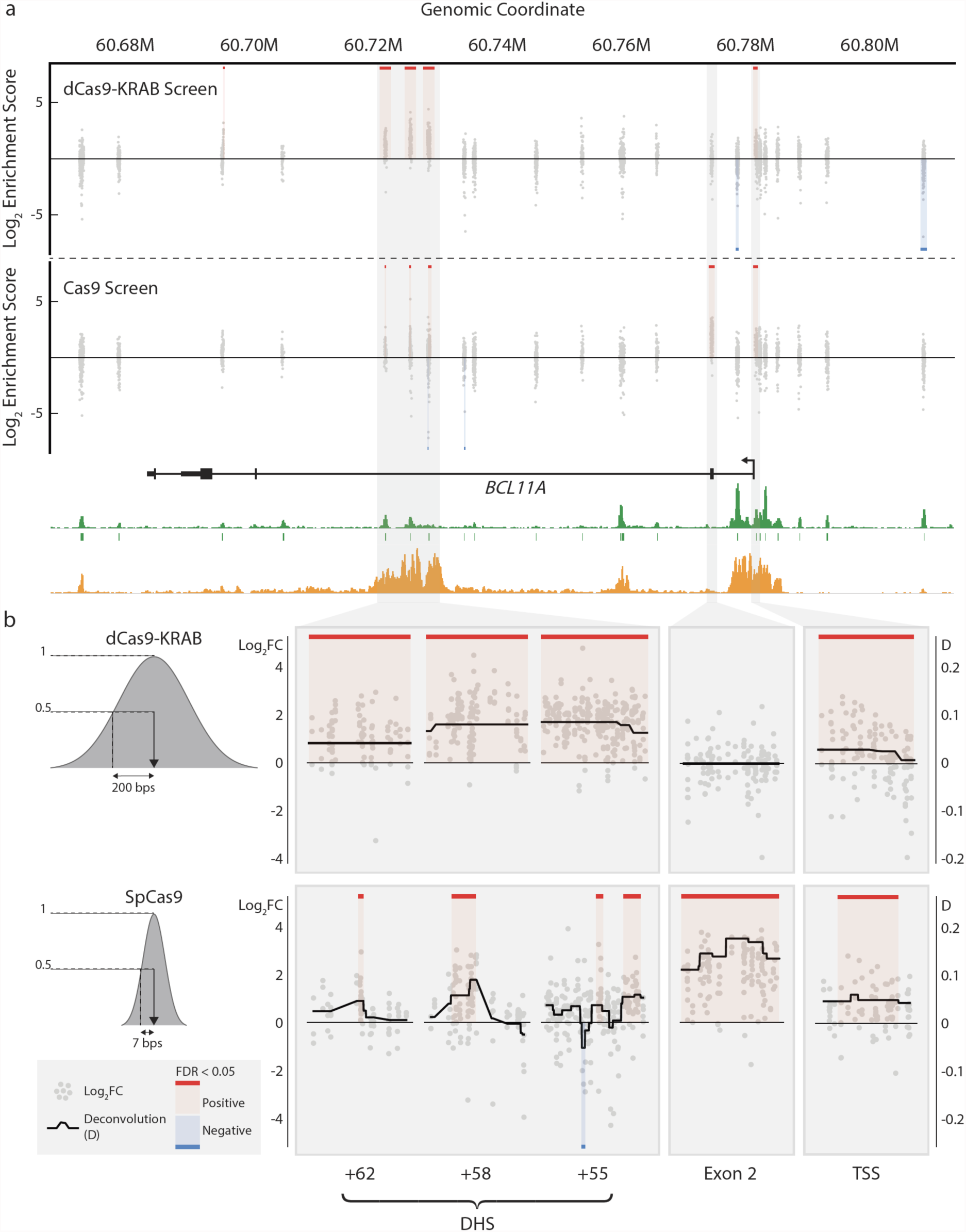
CRISPR-SURF analysis of *BCL11A* CRISPRi and CRISPR-Cas9 DHS tiling screens. **(a)** An overview of the *BCL11A* CRISPRi and CRISPR-Cas9 DHS tiling screens. **(b)** Shown are zoom-in panels of *BCL11A* exon 2 and common significant regions (FDR < 0.05) between the CRISPRi and CRISPR-Cas9 screens as determined by CRISPR-SURF.

CRISPR-SURF identified both previously known positive control regions in the *BCL11A* transcription start site (TSS) and exon 2 as significant. While the TSS was identified as a significant region in both the CRISPR-Cas9 and CRISPRi tiling screens, only the CRISPR-Cas9 screen returned exon 2 as significant, highlighting the necessity to introduce indel mutations rather than epigenetic modifications in the coding sequence to effectively disrupt *BCL11A* protein expression. In addition, three previously validated functional enhancers (DHS +55, +58, +62) regulating *BCL11A* were identified as significant regions in both the CRISPR-Cas9 and CRISPRi screens^1,11,12^. Importantly, the significant regions identified within these enhancers were narrower in the CRISPR-Cas9 screen compared to the CRISPRi screen, consistent with the narrower perturbation profiles of CRISPR-Cas9 indel mutations relative to CRISPRi epigenetic modifications. As expected, while the CRISPRi screen provided more power for detection (as evidenced by the greater number of supporting sgRNAs for each significant region), the CRISPR-Cas9 screen provided higher resolution of regulatory elements such as the identification of a significantly negative 41 bp region within DHS +55 that was not detected in the CRISPRi screen **(Figure 2b)**. In summary, CRISPR-SURF leverages the broad CRISPRi targeting window for efficient enhancer discovery and the narrow CRISPR-Cas9 perturbation profile for high-resolution mapping of critical elements within enhancers.

CRISPR-SURF is a user-friendly open-source software that can be used to design sgRNAs for tiling screens, to analyze new CRISPR tiling screen data, and to explore several pre-computed datasets at http://crisprsurf.pinellolab.org/ **(Supplementary Figure 8)**, or as a standalone command-line tool using Docker (https://github.com/pinellolab/CRISPR-SURF) **(Supplementary Note 5)**.

## Acknowledgements

We thank Chris Parmer for technical assistance with development of the CRISPR-SURF website. This work was supported by the National Institutes of Health (NIH) National Human Genome Research Institute (NHGRI) Career Development Award (R00 HG008399) (L.P.), the Defense Advanced Research Projects Agency (HR0011-17-2-0042) (L.P. and J.K.J.), NIH R03 DK109232 (D.E.B.), NIH DP2 OD022716 (D.E.B.), NIH P01 HL032262 (D.E.B.), Burroughs Wellcome Fund (D.E.B.), Doris Duke Charitable Foundation (D.E.B.), NIH R35 GM118158 (J.K.J.), NIH RM1 HG009490), the Desmond and Ann Heathwood Massachusetts General Hospital Research Scholar Award (J.K.J.), and the National Institute of Diabetes and Digestive and Kidney Diseases (NIDDK) Award (F30-DK103359) (M.C.C.).

## Author Contributions

J.Y.H. and L.P. conceived of and developed the CRISPR-SURF pipeline and web-application. M.A.C., M.C.C. and F.S. performed the *BCL11A* CRISPRi and CRISPR-Cas9 tiling screen experiments. D.P., C.P.F., J.M.E., E.S.L., R.F., K.C., S.H.O., and J.A.G. provided statistical and experimental expertise during the development of CRISPR-SURF. J.K.J., L.P., and D.E.B. oversaw the project and offered feedback and guidance. J.Y.H, L.P., D.E.B. and J.K.J wrote the manuscript with input from all authors.

## Data Availability Statement

Sequencing data from the *BCL11A* CRISPRi and Cas9 tiling screens is available at SRA accessions (submission pending approval).

## Code Availability Statement

The most-updated version of CRISPR-SURF as a command-line tool is available on GitHub at https://github.com/pinellolab/CRISPR-SURF, while the most recent version of the CRISPR-SURF desktop application can be found on DockerHub at https://hub.docker.com/r/pinellolab/crisprsurf/.

